# Plasma insulin-like growth-factor 1 (IGF-1) concentrations predict early life-history traits in a wild mammal

**DOI:** 10.1101/2025.06.02.656453

**Authors:** Sanjana Ravindran, Yolanda Corripio-Miyar, Joel L. Pick, Xavier Bal, Jill G. Pilkington, Josephine M. Pemberton, Daniel H. Nussey, Hannah Froy

## Abstract

1. The hormone insulin-like growth factor 1 (IGF-1) is a key player in the insulin/IGF-1 signaling (IIS) pathway. Extensive biogerontological research demonstrates that this evolutionarily conserved nutrient-sensing pathway plays a causal role in the regulation of growth, reproduction and longevity under laboratory conditions. However, its potential role as a mediator of adaptive life-history variation in highly variable natural environments remains unclear.
2. We measured IGF-1 concentrations in blood samples from approximately four-month-old wild Soay sheep lambs (n=669), collected over nine summers. We tested whether IGF-1 (i) was positively correlated with proxies of resource availability, (ii) was associated with morphological traits measured concurrently, and (iii) predicted subsequent fitness-related traits.
3. Plasma IGF-1 concentrations were higher in males compared to females, and positively correlated with measures of resource availability in both sexes. IGF-1 was lower in years of high population density when per capita food availability was reduced; in twin lambs who have fewer available resources compared to singletons; and in lambs born to young and old mothers, who have poor maternal provisioning compared to mothers of intermediate age.
4. Higher IGF-1 levels in summer were correlated with higher body mass, faster post-natal somatic growth and increased skeletal size, measured at the same time. These associations were independent of our proxies of resource availability.
5. Lambs with higher summer IGF-1 were more likely to survive their first winter and reproduce the following spring. The association between IGF-1 and reproduction was independent of our resource availability proxies, whereas the association with first-winter survival was not. The association between summer IGF-1 and reproduction was mediated by positive associations with summer body mass.
6. Our study reveals population-level phenotypic plasticity in circulating IGF-1, also finding IGF-1 to be positively associated with key morphological traits and positively predict fitness traits in early life. These findings highlight IGF-1 as a candidate physiological mechanism underpinning plastic responses to variation in food availability and influencing life-history traits in a wild mammal.

## Introduction

In fluctuating environments, individuals are selected to optimise allocation of limited resources between competing traits so that their fitness is maximised under current conditions. This generates phenotypic plasticity in life-history strategy, which is an important mechanism enabling organisms to cope with variable environments (Pigliucci., 2005). Although among-individual variation in life history has been the focus of considerable theoretical and empirical research, the physiological mechanisms underpinning adaptive life-history plasticity are poorly understood. The hormone Insulin-like Growth Factor 1 (IGF-1) has been proposed as a potential mediator of such effects, capable of integrating environmental cues and internal signals to coordinate a life-history response (Dantzer & Swanson., 2012).

IGF-1 is a key component of the evolutionarily conserved insulin/insulin-like growth factor 1 signaling (IIS) pathway, which senses nutrient availability and modulates energy metabolism and growth (Jones & Clemmons., 2005). This pathway is involved in the response to dietary restriction; the most robust environmental manipulation to extend lifespan in diverse taxa under laboratory conditions. It is therefore the subject of extensive biogerontological research which has revealed its role as an important regulator of ageing and lifespan across species (Mathew et al., 2017). In laboratory animal models, experimental downregulation of IIS signaling extends lifespan (Bartke et al., 2005; Berryman et al., 2008; Svensson et al., 2011; Kenyon et al., 1993). Exceptionally long-lived humans similarly exhibit lower circulating IGF-1 (Bonafe et al., 2003).

The link between nutrient availability and IGF-1 is extensively studied and robust (Regan et al., 2020; Clemmons & Underwood., 1991). In laboratory rodents and primates including humans, individuals on calorie-restricted diets had lower plasma IGF-1 concentrations (Mattison et al., 2017; Rahmani et al., 2019; Speakman & Mitchell., 2011). Previous work in laboratory rodents and domesticated vertebrates revealed IGF-1 to be crucial for somatic growth (Baker et al.,1993; Beccavin et al., 2001; Greer et al., 2011; Yakar et al., 2002). IGF-1 is also required for normal reproductive function in vertebrates; IGF-1 knockout mice were effectively infertile with smaller-sized reproductive organs (Baker et al., 1996; D’Ercole., 1999). In addition, experimental IGF-1 supplementation advanced puberty onset age in laboratory and domesticated mammals (Danilovich et al., 1999; Hiney et al., 1996).

Laboratory studies thus suggest that IGF-1 positively correlates with nutrient availability and has a causal influence on growth, reproduction and survival, however evidence from the wild is sparse (reviewed in Dantzer & Swanson., 2012; Lodjak & Verhulst., 2020). Experimental studies in wild bird, fish and reptile populations found plasma IGF-1 levels to be lower in food-restricted or fasting individuals compared to those fed *ad libitum* (Lodjak et al., 2023; Hack et al., 2018; Duncan et al., 2015). However, these experimentally manipulated diets may not reflect the fluctuating levels of food availability that wild populations inhabiting highly variable environments experience, leaving it unclear whether plasma IGF-1 shows similar patterns with food availability under natural conditions. Experimental and observational studies in wild vertebrates link higher IGF-1 to faster growth, larger body size and longer antlers (Crain et al.,1995; Lodjak et al., 2014, 2017, 2018; Lewin et al., 2017; Montoya et al., 2022; Tóth et al., 2022; Suttie et al., 1991; Ditchkoff et al., 2001). Whether these associations between IGF-1 and morphological traits persist after accounting for nutrient availability remains unclear. Even less is known about the link between IGF-1 and fitness traits in wild mammals – only one study in wild hyenas found juvenile females with elevated plasma IGF-1 levels to have earlier age at first parturition and reduced longevity after reproductive maturity (Lewin et al., 2017). Since heavier juveniles generally survive harsh conditions better and breed earlier, IGF-1 might influence fitness traits indirectly via body mass (Hamel et al., 2009; McMahon et al., 2015). Disentangling these associations demands field data that simultaneously track IGF-1, resource availability, body mass and life-history outcomes. Thus, despite strong laboratory evidence and promising avian and reptile studies in the wild, the role of IGF-1 on juvenile fitness traits remains virtually untested in free-ranging mammals.

In this study, we used plasma samples from juvenile wild Soay sheep collected over nine summers to test hypotheses about IGF-1 under natural conditions. We hypothesized that levels of circulating IGF-1 would: (i) vary in response to resource availability, being higher when population densities are low and per capita food availability is consequently high, in singleton lambs that do not compete for maternal resources, and in lambs born to prime-aged mothers who are better able to provision their offspring; (ii) correlate positively with morphological traits including post-natal growth, structural size, body condition and secondary sexual traits (horn length); and (iii) positively predict subsequent life-history traits including first-year reproduction and overwinter survival. To understand whether the associations between IGF-1 and the morphological and life-history traits were independent of our proxies of resource availability and/or mediated by body mass, we estimated the effects of IGF-1 conditional on these predictors. By doing so, we could disentangle the intricate relationships between IGF-1, resource availability proxies, body mass and life-history traits, enabling us to understand the role of IGF-1 in underpinning life-history variation in the wild (Figure S1).

## Materials and Methods

### Study system and data collection

Since 1985, the Soay sheep resident in the Village Bay area of Hirta (around 30% of the island’s population) have been subject to individual-based monitoring (Clutton-Brock & Pemberton, 2004). Population density fluctuates among-years, with most sheep mortality occurring February–April and biased towards lambs and males. Regular censusing throughout the year along with mortality searching allows accurate estimation of death dates. Both males and females are sexually mature in their first year. The Soay sheep are polygynous with intense male–male competition for access to females during the rut in October–November. Successful pregnancies result in ewes giving birth in March–May, although still-births also occur. The pregnancy status of females that die over winter is assigned through postmortem assessment. The accurate assignment of both parents to offspring is performed using genome-wide SNP markers and the pedigree inference program SEQUOIA (Huisman, 2017). Lambs are fully weaned by the end of autumn (Clutton-Brock & Pemberton, 2004). Most females produce singleton offspring, but depending on environmental conditions, up to 25% can give birth to twins in a given year (Clutton-Brock et al, 1991).

Around 95% of the lambs born in the study area are caught, weighed and uniquely tagged for identification within a few days of their birth (Robertson et al., 1992). Each August, a fieldwork team visits Hirta and captures as many of the animals resident in the Village Bay area as possible (usually 50-60% are caught within a two-week period). Temporary traps are constructed across the study area and at capture, blood samples and phenotypic measurements such as body weight, foreleg length, and horn length are taken from all individuals (Clutton-Brock & Pemberton., 2004). Blood samples are collected in lithium heparin tubes and kept at 4°C until processing, usually within 24 hours of sampling. The tubes are spun at 1,000g for 10 minutes. The plasma layer is transferred to 1.5 ml Eppendorf tubes and then stored at -20°C.

### Insulin-like Growth Factor 1 Enzyme-Linked Immunosorbent Assays (IGF-1 ELISAs)

Our initial dataset was comprised of lambs born over 9 breeding seasons, between 2014 and 2022 (n=683 individuals). We measured IGF-1 concentrations of these approximately 4-month-old lambs from archived plasma samples. All 683 samples were assigned randomly to different ELISA plates (n=18) and run in duplicate. We used a commercially available ovine enzyme-linked immunoassay kit (Sheep IGF-1 ELISA Kit, abx576572; Abbexa Ltd) following manufacturer’s instructions. Briefly, a 60 ng/ml standard solution was serially diluted six times to obtain a standard curve. Plasma samples were diluted to 1:25 in phosphate-buffered saline solution (1X PBS with pH 7.2; Gibco) to ensure our measurements fell within the range of the kit standard curve (0.94 ng/ml to 60 ng/ml). Two blank wells containing the diluent buffer were also included as negative controls. After aliquoting 100µl of standard and blank solutions into the ELISA plate wells, 100µl of the diluted samples was added in duplicate. The plates were then sealed and incubated at 37°C for 1 hour. After discarding the liquid, 100µl of Detection Reagent A (primary antibody) was added to all wells. Plates were once again sealed and incubated for 1 hour at 37°C. Plates were then washed 3 times with 1X Wash Buffer using an automated microplate washer (Skan Washer 400; Molecular Devices). This was followed by the addition of Detection Reagent B (secondary antibody) and incubation for 30 minutes at 37°C. Following another wash step (5 times with 1X Wash Buffer), 90 µl of TMB substrate was added to the wells and plates were incubated in the dark for 8 minutes following which 50 µl of stop solution was added. Using a microplate reader (FLUOstar Omega; BMG Labtech), the optical density of samples was read at 450 nm. The average concentration of duplicate samples was estimated as the plasma IGF-1 concentration for the individual. Samples were re-run if the difference in optical density (OD) between duplicates was > 0.2 (n=2).

There were 14 samples (out of 683 i.e. 2.05% of full dataset) for which the intra-assay coefficient of variation was >=11% and were excluded from further analyses. To determine inter-plate repeatability, we ran one ELISA plate (n=40 samples in duplicate) on consecutive days with samples placed in the same location on both plates. We then fitted a linear mixed effect model including sample identity as a random intercept; repeatability was estimated as the proportion of variance explained by sample identity over the total variance. This inter-plate technical repeatability of the ELISA assays was estimated as 94.524%. We also tested for effects of blood sample collection time and freezer storage time of plasma samples on IGF-1 concentrations (Supplementary Methods & Results).

### Definitions of variables

#### Proxies of resource availability

We considered population density, litter size and maternal age at birth as indirect measures of resource availability. In this population, food is a limiting factor with increased mortality rates during winter when population density is high. Twins are physically smaller at birth and throughout early life compared to singleton lambs. In addition, lambs born to younger and older mothers have lower body mass at birth and poorer survival outcomes in their first year, compared to those born to mothers in their prime (Clutton-Brock & Pemberton., 2004).

Annual population density was estimated as the total number of individuals in the study area alive on October 1^st^ in the year of measurement (which is highly correlated with population density in late spring and summer, given the very low mortality over this period). Litter size was a categorical variable with individuals classified as being born either a singleton or twin. Maternal age at birth was the age of the mother (in years) at offspring birth. We used the average maternal age in the dataset (5.456 years) for those observations where birth year of the mother was unknown (n=44 lambs).

#### Morphological Traits

Morphological trait data used in this study included a) August body mass – the weight of an individual at capture in August, to the nearest 0.1 kilogram (kg) (n=666 individuals); b) Post-natal growth – defined as August body mass of an individual controlling for birth weight (in kg) and age (in days) (n=545 observations); c) August foreleg length – a proxy for skeletal size, measured as the length of the metacarpal with hoof and knee joint bent away from it, to the nearest millimeter (mm) (n=660 observations); d) August horn length – for normal-horned individuals, measured along the outer curvature from the base of the horn to the tip, to the nearest mm (n=361 individuals; IGF-1 did not vary depending on individual horn type, see Supplementary Methods & Results - Table S3).

#### Life-History Traits

We defined first-year reproduction as a binary outcome with individuals who produced an offspring in their first year (determined using the pedigree) assigned ‘1’ and those who did not produce offspring assigned ‘0’. All female lambs were included in this analysis (n=337) since reproductive status could be determined even after death by *postmortem*. For male lambs, only those individuals seen on at least one rut census were included (n=319). In the complete dataset (n=656 individuals), 68 individuals had offspring that were still-births or foetuses that died *in utero* (10.37%) while 89 individuals produced live offspring (13.57%). There were 134 females and 23 males assigned ‘1’ for reproduction (including foetuses, still births and live offspring) while 499 individuals were assigned ‘0’. Like reproduction, first-year overwinter survival was a binary response with individuals that survived to June 1^st^ of the following year assigned ‘1’ and those who died before June 1^st^ the following year assigned ‘0’ (n=609 individuals).

### Statistical Analyses and Predictions

#### Do proxies of resource availability predict summer plasma IGF-1 concentrations in lambs?

We first investigated whether our proxies of resource availability predicted summer plasma IGF-1 concentrations (n=669). We ran a linear mixed effects model with population density (linear term), litter size (two-level factor) and maternal age at birth (linear and quadratic terms) as predictor variables and IGF-1 concentration as the response (Figure S2). Sex was included as a categorical predictor to account for differences in levels of circulating IGF-1 between males and females. To account for assay-related variation, ELISA plate number and run date were included as random effects. We also included random effects for sample year and maternal identity to account for repeated measures across years and across mothers. These models were run using the ‘*rstanarm’* package in R using default weakly informative priors for all parameters. P-values for fixed effects were estimated using the ‘*parameters’* R package and represent the probability of direction (strongly correlated with frequentist p-value) (Makowski et al., 2019). Conditional and marginal R^2^ were estimated using the ‘*performance’* R package.

#### Are plasma IGF-1 levels associated with morphological traits?

We ran separate linear mixed-effects models with body mass, post-natal growth, skeletal size and horn length (only in normal-horned individuals) as response variables. We fitted IGF-1 as a continuous fixed effect, and sex as a categorical predictor to account for differences in mean trait values between males and females. Random effects of maternal identity and year were also fitted. These models were fitted using an errors-in-variables modelling approach using ‘*cmdstanr’* R package (see below for further details).

Since variation in both IGF-1 and morphological traits may be driven by variation in resource availability, we re-ran the above models including population density (linear term), litter size (two-level factor) and maternal age (linear and quadratic terms) as additional predictors to investigate the independent associations between IGF-1 and morphological traits, controlling for our proxies of resource availability. We also re-ran the foreleg and horn length models including body mass as an additional covariate to understand whether the associations with IGF-1 were independent of summer body mass (Table S6). All continuous predictors were standardized to mean = 0 and standard deviation = 1 before including in the model.

#### Do plasma IGF-1 levels predict first-year reproduction and survival over the first winter?

We ran separate generalized linear mixed-effects models of first-year reproduction and overwinter survival, fitting them as binary response variables. IGF-1 was fitted as a continuous predictor and sex was included as a categorical predictor. Maternal identity and year were included as random effects. These models were again fitted using an errors-in-variables modelling approach using ‘*cmdstanr’* R package (see below for further details).

To understand if IGF-1 levels predicted first-year reproduction and survival independent of resource availability, we re-ran the above models including our proxies of resource availability as additional fixed effects. Since body mass could mediate the association between IGF-1 and life-history traits (Figure S1), we also re-ran this model including summer body mass as an additional predictor. All continuous predictors were standardized to mean = 0 and standard deviation = 1 before including in the model. Sample sizes for all analyses and across different categories are described in Table S1 and S2.

#### Errors-in-variables modelling approach

Measurement error in predictor variables leads to a general attenuation of effects. Given that almost half of the variation in IGF-1 was due to lab-related measurement error (48.445 % of variation explained by ELISA plate number and run date; Table 1), we employed an errors-in-variables modelling approach when investigating whether IGF-1 predicted morphological and life-history traits. This approach accounts for measurement error in the predictors and reduces bias in the estimated effects (NB – When IGF-1 was fitted as the response variable in earlier models, these sources of measurement error were accounted for as random intercept terms and so, an errors-in-variables modelling approach was not employed there).

**Table 1.**
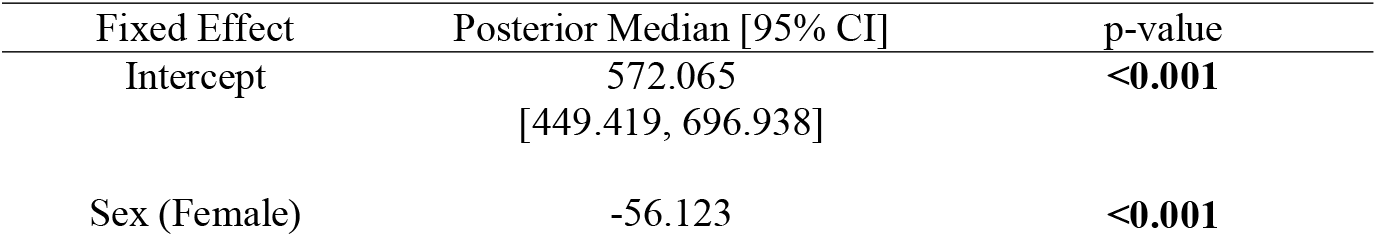
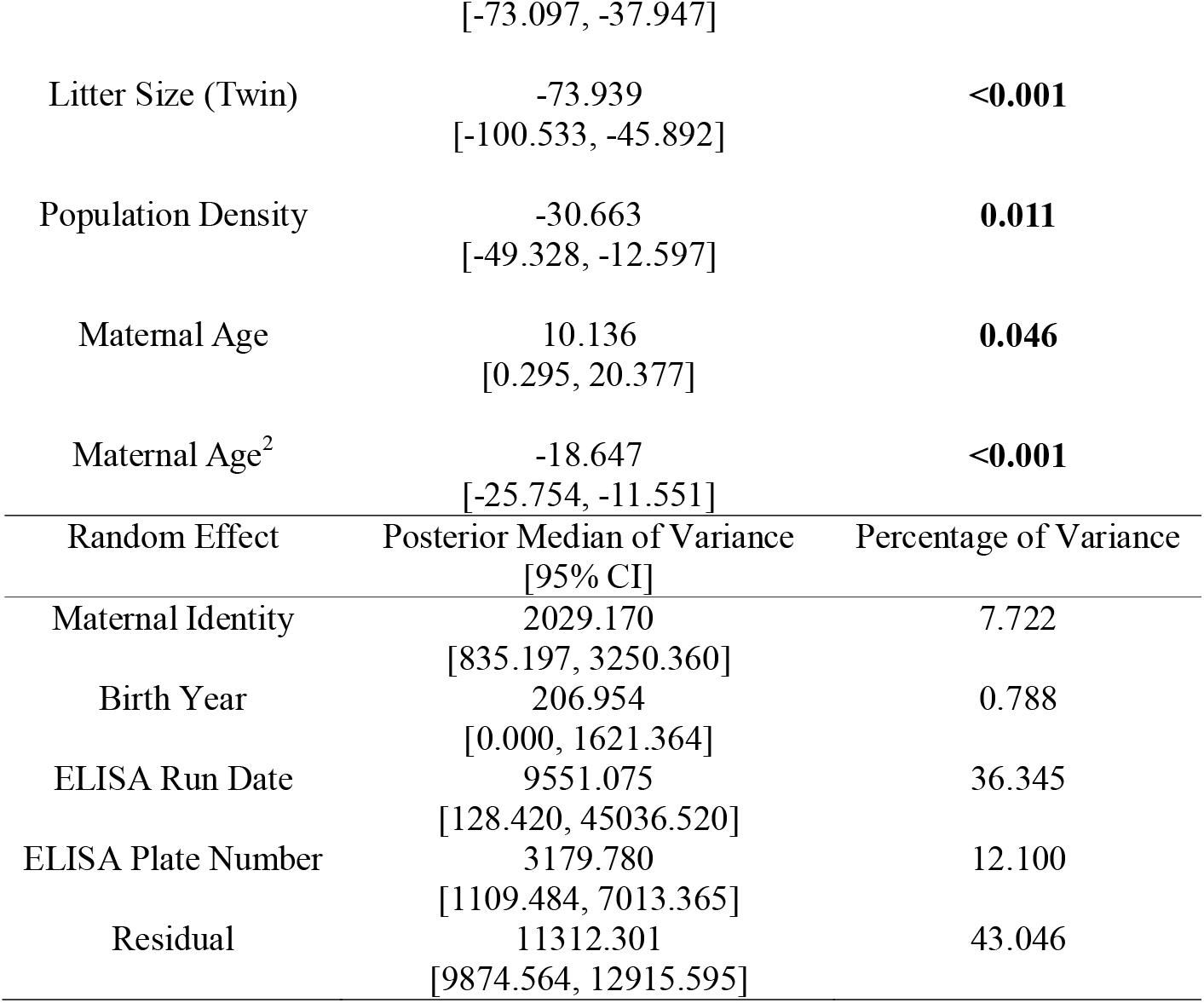
Result from linear mixed effects model investigating variation in IGF-1 with sex, population density, litter size and maternal age at birth (n=669 observations; population density and maternal age at birth were standardized to mean=0 and standard deviation=1; reference levels for the factors were sex: male, litter size: singleton)

In this approach, we assumed that the ‘corrected IGF-1’ value is a latent variable that can be expressed as –

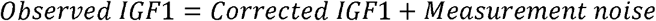

Corrected IGF-1 values can be estimated according to the above equation and inputted in our subsequent models investigating if *IGF*1_*Corrected*_ predicted morphological and life-history traits.

We jointly modelled the measurement error in ‘observed IGF-1’ and whether ‘corrected IGF-1’ predicted variation in morphological and life-history traits using Stan (implemented using the ‘*cmdstanr’* R package version 0.8.1). Prior to model fitting, observed IGF-1 was scaled to have a mean of 0 and a standard deviation of 1. We specified the likelihood for ‘observed IGF-1’ for an individual *i* as normally distributed with a mean of 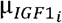 and a standard deviation *σ*_*IGF*1_,

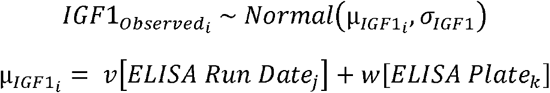

Where *IGF*1_*obs*_ is the observed IGF-1, *v* is the intercept varying by ELISA run date *j* ∈ {1,2,3,4} and *w* is the intercept varying by ELISA plate *k* ∈ {1,2,3, …18}.

We defined the relationship between ‘observed IGF-1’ and ‘corrected IGF-1’ values as follows –

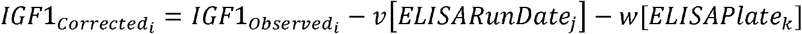

These ‘corrected IGF-1’ parameter estimates were fitted as a predictor variable in models where either morphological or life-history traits were the dependent variables:

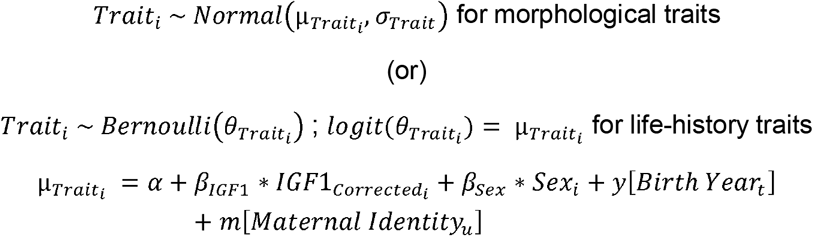

Where a trait value for an individual i is either normally distributed with a mean of μ_*Trait*_ and a standard deviation of *σ*_*Trait*_ (for morphological traits) or specified as a Bernoulli distribution with canonical logit link (for life-history traits) where 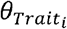 is the probability of success. The expected trait value 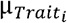 can be expressed as a regression equation with an intercept *α, β*_GFl1_ as the slope for 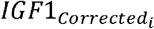 and *β*_*sex*_ as the slope for *sex*_*i*_ (coded as a dummy variable with females assigned ‘1’ and males assigned ‘0’). y is the intercept varying by birth year *t* ∈ {1,2, … 9} and m is the intercept varying by maternal identity *u* ∈ {1,2, … 306}.

The weakly-informative prior for the error term, *σ*_*e*_ was defined as a Cauchy distribution with location parameter (mean) = 0 and scale parameter (standard deviation) = 2.5. The random intercepts v and w are assumed to be normally distributed with a mean of 0 and standard deviations *σ*_*v*_ and *σ*_*w*_ respectively, where *σ*_*v*_, *σ*_*w*_ were also specified as Cauchy distributions (mean=0, sd=2.5).We used non-centered parameterization to define priors for the random intercepts *y* and *m*, assumed to be normally distributed with a mean of 0 and standard deviations *σ*_*y*_ and *σ*_*m*_ respectively, specified as Cauchy distributions (mean=0, sd=2.5). Default priors were used for all other parameters in our models. We ran four Markov chains in parallel with 2000 iterations per chain (1000 for warm-up and 1000 for sampling). We checked model diagnostics by evaluating the Rhat parameter equals 1 and the effective sample size > 400 in the bulk and tail of the posterior distribution for all parameter estimates. We also performed posterior checks by comparing distributions of 200 posterior draws to the observed data (Figure S4). All analyses were run using R version 4.4.1 (2024-06-14) in RStudio (2024.09.0+375).

## Results

### Proxies of resource availability predict summer IGF-1 concentrations in lambs

We observed lower plasma IGF-1 concentrations in females compared to males (Figures 1A & S3, Table 1). Plasma IGF-1 levels were negatively associated with population density (Figure 1C, Table 1). In addition, being born as a twin was associated with decreased plasma IGF-1 levels in the summer relative to being a singleton (Figure 1B, Table 1). A quadratic relationship between IGF-1 and maternal age was observed; lambs born to younger and older mothers had reduced IGF-1 levels compared to those born to prime-age mothers (Figure 1D, Table 1). The conditional R^2^ (variance explained by both fixed and random effects) was estimated as 0.552 (95% CI: 0.501 – 0.598) while the marginal R^2^ (variance explained by fixed effects alone) was 0.121 (0.075 – 0.167).

**Figure 1.**
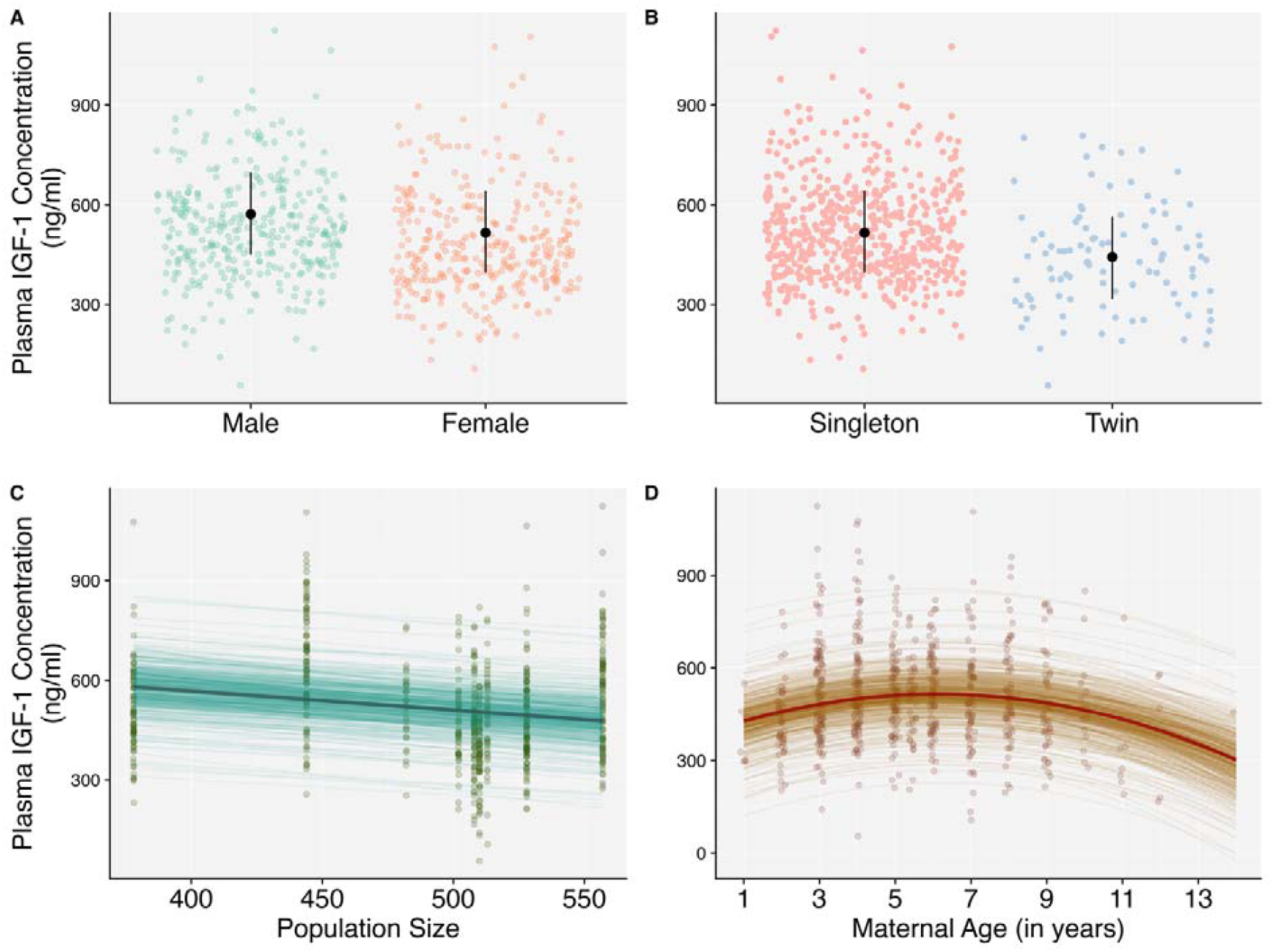
Association between plasma IGF-1 levels and A) Sex, B) Litter size at birth, C) Population density and D) Maternal age at birth in Soay sheep lambs (n=669 observations; Table 1). Black points and error bars represent posterior median and 95% CIs in A and B. Posterior median and 500 draws from joint posterior distribution represented in C and D by dark- and light-shaded lines respectively. Coloured points represent raw data.

### Plasma IGF-1 levels are positively correlated with body mass, skeletal size and postnatal growth

Increased levels of plasma IGF-1 were associated with heavier body mass and larger structural size in lambs (Table S6A, S6C). After accounting for variation in birth weight and lamb age at measurement (in days), we still observed a positive association between plasma IGF-1 levels and summer body mass (Table S6B), suggesting that higher plasma IGF-1 levels are associated with faster post-natal growth. We also observed a positive correlation between IGF-1 levels and horn length in normal-horned individuals (Table S6D). These positive associations between IGF-1 and all morphological traits were present after controlling for our proxies of resource availability (Figure 2; Table S6). Additionally, we observed independent positive associations between IGF-1 and both structural size and body mass (Table S6). However, the association between IGF-1 and horn length was no longer statistically significant after accounting for summer body mass (Table S6D).

**Figure 2.**
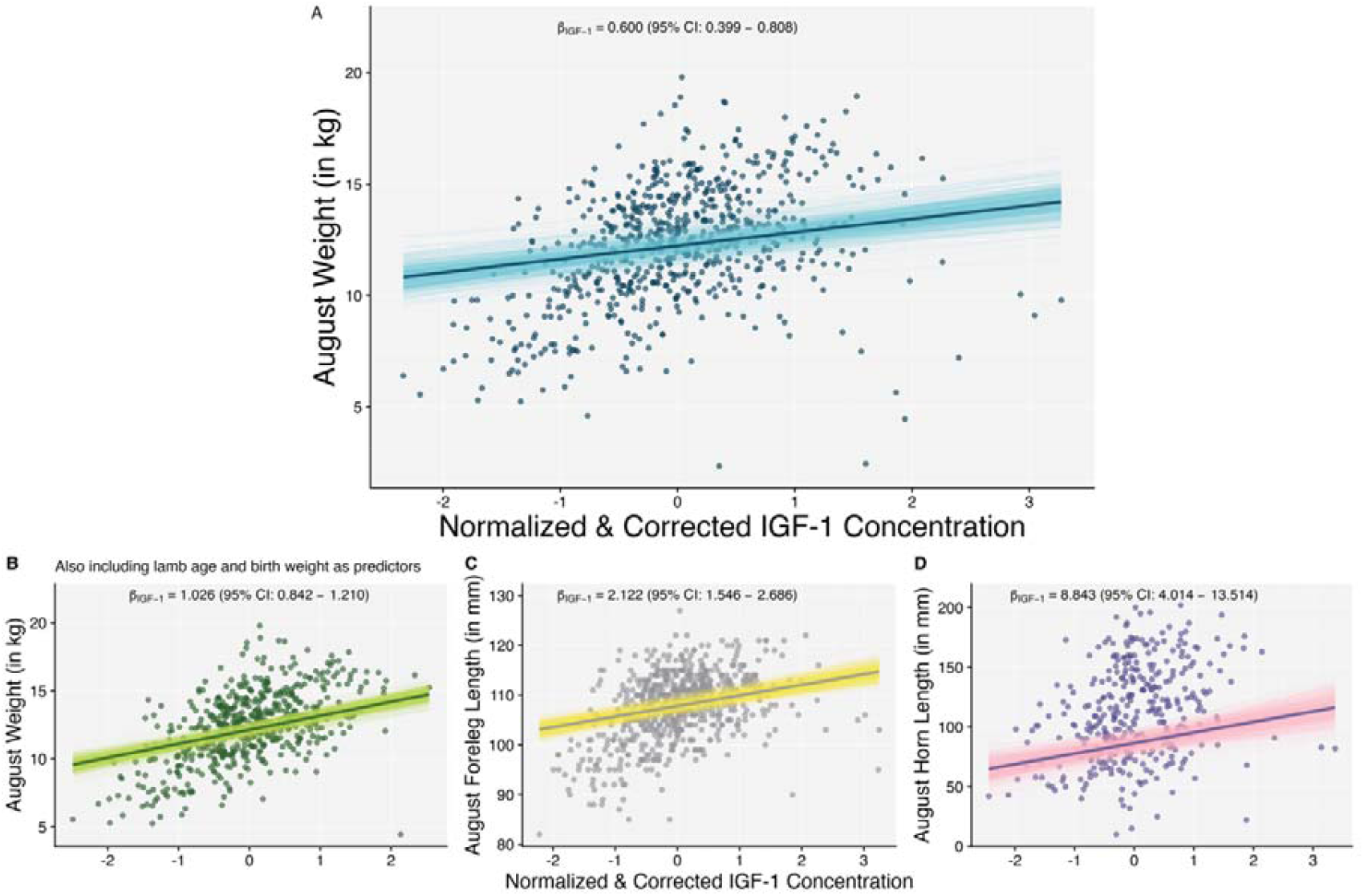
Associations between plasma IGF-1 levels and A) August body mass (n=666); B) Post-natal growth (n=545); C) August foreleg length (n=660); D) August horn length (n=361) in Soay lambs, independent of our proxies of resource availability (population density, sex, litter size and maternal age at birth; Table S6). The slope is predicted with fixed effects set as follows: female sex, singleton litter, average population density and average maternal age at birth. The dark line represents the median of the posterior distribution and the lightly shaded lines are 500 draws from the posterior distribution. Points represent raw data (*IGF*1_*Corrected*_ estimated from *IGF*1_*Observed*_ which was standardized to mean=0 and standard deviation =1 prior to model fitting).

### Plasma IGF-1 concentrations predict first-year survival and reproduction

We found lambs with higher plasma IGF-1 concentrations in the summer were more likely to reproduce in their first year (Table S7A). This association was present after controlling for resource availability proxies (Figure 3A; Table 2). However, after accounting for summer body mass, there was no statistical support for plasma IGF-1 levels positively predicting reproduction (Figure 3B, Table S7A). This is consistent with the association between IGF-1 and reproduction being mediated by summer body mass.

**Figure 3.**
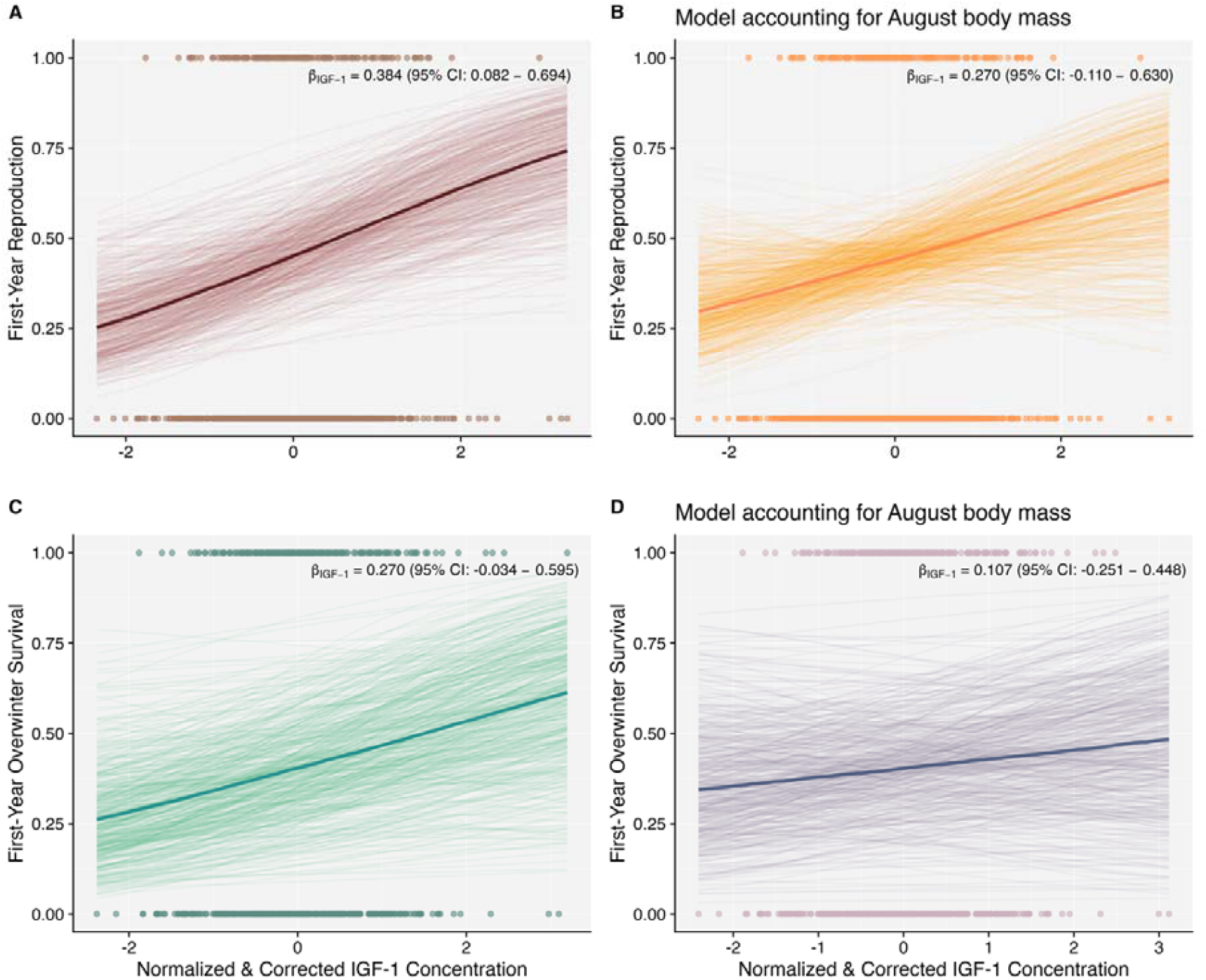
Associations between plasma IGF-1 levels and first-year reproduction (n=656) and first-year overwinter survival probability (n=609) in lambs, independent of resource availability proxies (A and C) or both resource availability and summer body mass (B and D) (Table S7). The slope is predicted with fixed effects set as follows: female sex, singleton litter, average population density and average maternal age at birth. The dark line represents the median of the posterior distribution and the lightly shaded lines are 500 draws from the posterior distribution. Points represent raw data (*IGF*1_*Corrected*_ estimated from *IGF*1_*Observed*_ which was standardized to mean=0 and standard deviation =1 prior to model fitting).

**Table 2.** Results from (generalized) linear mixed effects models investigating the associations between circulating IGF-1 levels and summer body mass (n=666 observations), first-year reproduction (n=656 observations) and first-year overwinter survival probability (n=607 observations) in lambs. Reference levels for the factors were sex: male, litter size: singleton.

|  | Summer Body Mass | First-Year Reproduction | First-Year Overwinter Survival |
| --- | --- | --- | --- |
|  | 13.978 | -2.596 | -0.908 |
| Intercept | (13.141 – 14.849) | (-3.467 – -1.764) | (-2.282 – 0.296) |
|  | **0.600** | **0.384** | 0.270 |
| IGF-1 | **(0.399 – 0.808)** | **(0.082 – 0.694)** | (-0.034 – 0.595) |
|  | **-1.096** | **2.661** | **0.986** |
| Sex (Female) | **(-1.404 – -0.789)** | **(2.113 – 3.277)** | **(0.528 – 1.466)** |
|  | **-2.983** | **-1.399** | **-0.998** |
| Litter Size (Twin) | **(-3.498 – -2.497)** | **(-2.241 – -0.607)** | **(-1.817 – -0.287)** |
|  | 0.063 | -0.320 | -0.660 |
| Population Density | (-0.827 – 0.959) | (-1.125 – 0.439) | (-1.936 – 0.595) |
|  | **0.475** | 0.112 | -0.013 |
| Maternal Age | **(0.297 – 0.651)** | (-0.120 – 0.353) | (-0.283 – 0.241) |
|  | **-0.652** | **-0.262** | **-0.456** |
| Maternal Age<sup>2</sup> | **(-0.778 – -0.526)** | **(-0.476 – -0.064)** | **(-0.706 – -0.231)** |
| Variances |  |  |  |
|  | 1.411 | 0.120 | 1.033 |
| Maternal Identity | (0.915 – 2.029) | (0.000 – 0.809) | (0.262 – 2.468) |
|  | 1.133 | 0.792 | 2.195 |
| Birth Year | (0.382 – 4.376) | (0.172 – 3.711) | (0.729 – 8.578) |
|  | 2.875 |  |  |
| Residual | (2.502 – 3.360) |  |  |

Lambs with higher plasma IGF-1 had higher survival probabilities over their first winter (Table S7B). The slope between plasma IGF-1 and overwinter survival was positive, however the 95% CIs overlapped zero in models where we controlled for resource availability proxies, body mass or both (Tables 2, S7B; Figures 3C, 3D).

## Discussion

We measured levels of circulating IGF-1 in plasma samples collected from 669 juvenile wild Soay sheep over nine summers. IGF-1 was positively correlated with proxies of resource availability, being higher in years of low population density, and in singleton lambs born to prime-aged mothers. IGF-1 was associated with multiple morphological traits, positively correlating with body mass, post-natal growth and skeletal size. Higher plasma IGF-1 predicted both first-year reproduction and overwinter survival. However, we found no statistical support for the IGF-1–survival association after accounting for either resource availability proxies or summer body mass. IGF-1 predicted first-year reproduction independent of resource availability, but this association was mediated by summer body mass. Our findings reveal population-level phenotypic plasticity in plasma IGF-1 in response to variation in resource availability, and support for a potential role for plasma IGF-1 as a physiological marker of total resource budget and a regulator of life-history plasticity in the wild.

Circulating levels of IGF-1 varied predictably with our proxies of resource availability, including measures of both the maternal and external environment. Soay lambs rely largely on their mother’s milk for their nutritional needs until they become fully weaned in late summer (Clutton-Brock and Pemberton., 2004). In years when population density is high, forage biomass and primary productivity decline, intensifying competition for food (Robertson et al., 1992). Lambs born in such years could thus experience reduced nutrient levels due to both their mothers having less to eat and when the lambs become independent, they have fewer resources available to feed. Previous experimental studies in wild bird and fish populations similarly found plasma IGF-1 concentrations to be reduced in fasting or food-restricted individuals (Schew et al., 1996; Gabillard et al., 2006; Uchida et al., 2003; Wilkinson et al., 2006). Resource limitation effects on juvenile IGF-1 can also act via maternal traits. In most iteroparous mammal populations, individuals prioritize allocation of resources towards survival over reproduction when resources are scarce (Therrien et al., 2008). As a result, reduced maternal allocation of resources to offspring during gestation and lactation could lower juvenile IGF-1 levels. Young mothers are also resource-poor as they are still growing themselves while older mothers are in poor physiological condition and so are less able to meet the resource demands of gestation and lactation (Hamel et al., 2012). Even within litters or broods, sibling competition for shared maternal resources affected circulating IGF-1 levels – great tits and jackdaws nestlings from reduced broods had higher IGF-1 than those in enlarged or control broods (Lodjak et al., 2014; Lodjak et al., 2023 but see Ridenour et al., 2023). We also found sex differences in IGF-1 levels, with elevated IGF-1 in male compared to female Soay sheep. In Soays, this likely reflects greater food intake or faster growth rates of male lambs in this sexually dimorphic species (Clutton-Brock & Pemberton 2004). This result of high IGF-1 levels in the larger sex is consistent with findings in domestic animals and wild reptiles (Reda et al., 2024; Baéza et al., 2001; Meter et al., 2022; Roberts et al., 1990). Overall, our results add to a growing body of evidence suggesting that levels of circulating IGF-1 vary plastically in response to resource availability in the wild.

We found elevated levels of plasma IGF-1 in heavier and larger 4-month-old Soay sheep. Faster-growing individuals also had high IGF-1 levels. This finding is consistent with evidence from laboratory studies that circulating IGF-1 promotes body growth and bone development in vertebrates. The IGF-1/Akt/mTOR pathway is the key mediator of cell proliferation and survival across diverse tissues (Hakuno & Takahashi., 2018). IGF-1 is also part of the growth hormone (GH)/IGF1 axis mediating the effects of GH on somatic growth (Laron., 2001). Our finding is also in line with results from previous observational and experimental studies in wild vertebrates (Ditchkoff et al., 2001; Lodjak et al., 2017; Tóth et al., 2022). A positive association between plasma IGF-1 and tarsus length - a proxy for structural body size in birds - was also observed in two free-living passerine populations (Lodjak et al., 2014, 2017). Crucially, we observed consistent positive associations between IGF-1 and all morphological traits even after controlling for our proxies of resource availability. This indicates that IGF-1 captures morphological trait variation that is not fully explained by our resource proxies. It is likely though that there is additional variation in resource availability that is not fully captured by our proxies. For instance, differences in vegetation quality between years or within-year variation in space-use behaviour, home-range quality and local population density may be associated with both IGF-1 and morphological traits (Crawley et al., 2021; Wiersma et al., 2023). Our results suggest that IGF-1 could act as a physiological marker of body size and post-natal somatic growth and may play a role in mediating resource allocation decisions towards key morphological traits in wild mammals.

Understanding the genetic correlation between IGF-1 and body size in future studies can help reveal the pleiotropic nature of this hormonal pathway in directing organismal development and shaping individual life-histories.

The influence of plasma IGF-1 on short-term survival prospects across both laboratory and wild environments is unclear. We found variation in summer plasma IGF-1 to predict survival over the lamb’s first winter. However, this association was not independent of resource availability proxies and body mass, suggesting that IGF-1 is not a proximate driver of juvenile first-year survival in this population. In contrast to our results, in wild female spotted hyenas, plasma IGF-1 levels did not predict survival to reproductive maturity but instead, predicted adult survival (Lewin et al., 2017). Reduced IIS activity consistently predicts longer lifespan in laboratory studies across diverse species (reviewed in Mathew et al., 2017). Comparative studies in wild mammals and birds similarly found higher IGF-1 levels to be associated with fast life-histories (Lodjak et al., 2018; Swanson & Dantzer., 2014). Our study could not investigate associations between plasma IGF-1 and adult lifespan since many individuals that survived their first winter are currently alive and have not yet expressed their full lifespan. Repeated measures of plasma IGF-1 concentrations of individuals across their lifespan will be essential to resolve whether IGF-1 actively shapes survival trajectories or simply reflects variation in resource levels or body condition. This will illuminate how IGF-1 may regulate lifespan and the natural ageing process in wild populations.

Higher summer IGF-1 levels predicted increased probability of first-year reproduction in Soay lambs. Although independent of variation in resource availability proxies, this association between IGF-1 and reproduction was explained by variation in summer body mass – individuals with higher plasma IGF-1 concentrations were heavier and so, more likely to reproduce in their first year. IGF-1 is part of the hypothalamic-pituitary-somatotropic (HPS) axis which interacts strongly with the hypothalamic-pituitary-gonadal (HPG) axis, the latter being the key facilitator of reproduction and sexual behaviours in vertebrates (Milman et al., 2016; Sower et al., 2009). It is likely that fertility is mainly regulated by hormones involved in the HPG axes with IGF-1 contributing to this primarily via its influence on body size. Similar to our findings, previous studies in laboratory and wild populations have suggested a size-mediated effect of IGF-1 on reproductive traits (Kroonsberg et al., 1989; Lodjak et al., 2018). In contrast, a study in a wild population of juvenile female hyenas found plasma IGF-1, but not body mass, better predicted age at first parturition although sample sizes were relatively small (n=37; Lewin et al., 2016). Positive links between plasma IGF-1 and reproductive traits have been previously reported in laboratory rodents, cattle, wild reptiles and fish (D’Ercole., 1999; Daftary & Gore., 2005; Patton et al., 2007; Sparkman et al., 2010; Campbell et al., 2006). But it is unclear in these studies whether resource availability confounds the association between IGF-1 and reproduction, or whether the observed IGF-1–reproduction association is mediated by effects of IGF-1 on body mass. Importantly, our reproductive metric included pregnancies that resulted in either foetal loss, still-births or successful births, which suggests that IGF-1-driven somatic growth confers advantages already at conception and very early gestation. Similar positive associations between plasma IGF-1 and early fertility traits like first-service conception rates and pre-implantation embryo development have been reported in dairy cows (Patton et al., 2007; reviewed in Velazquez et al., 2005). Our findings provide support for plasma IGF-1 as a potentially exciting mechanistic link between resource availability, juvenile growth and first-year reproduction in Soay sheep, capable of shaping life-history variation in wild vertebrates.

## Supporting information

Supplementary Methods and Results

## Acknowledgements

Many thanks to all project members and volunteers who have all contributed immensely to the St. Kilda Soay Sheep Project over many years, particularly Ian Stevenson for managing the Soay sheep database. We are grateful to the National Trust for Scotland for permission to work on St Kilda and QinetiQ and Kilda Cruises for logistical support in the field. We thank the National Environment Research Council for funding the long-term project and the Royal Society for funding the research conducted in this study. Figure 1 in the supplementary file was created using BioRender.

## Author Contributions

SR and HF conceived the study; SR performed the laboratory work and statistical analyses with support from YCM and JLP respectively; JMP led pedigree reconstruction; DHN, JMP, JGP, XB are involved in management of the long-term study and fieldwork; SR wrote the first draft of the manuscript with input from HF; all authors contributed to final revisions and editing of the manuscript.

