## Supplementary Methods and Results for "Plasma insulin-like growth-factor 1 (IGF-1) concentrations predict early life-history traits in a wild mammal"

The Soays exhibit three different horn types: normal, scurred (misshapen horns) and polled (no horn) (Robinson et al., 2006). To understand if IGF-1 predicted horn length in normal-horned individuals, we first ran a model testing whether IGF-1 levels differed across the three horn-types (n=667 observations). This model included sex (factor), population size (continuous), litter size (factor) and maternal age at birth (linear and quadratic terms) as additional predictors. Population density and maternal age were scaled to mean=0 and standard deviation=1, prior to model fitting. Random effects of ELISA plate number, ELISA run date, individual identity and birth year were also fitted. We found no differences in mean IGF-1 levels according to horn-type (Figure S5, Table S3).

To evaluate whether sample storage time influenced plasma IGF-1 concentrations, we re-ran the IGF-1 model including storage time (in years) as an additional continuous covariate (n=669 observations). Here, IGF-1 was the response variable and fixed effects of population density (continuous), sex (factor), litter size (factor) and maternal age at birth (linear and quadratic terms) were fitted. All continuous predictors were z-standardized prior to model fitting. Random effects of ELISA plate number, ELISA run date, maternal identity and birth year were also included. We found a small but statistically significant positive association between storage time and plasma IGF-1 (Figure S6; slope=17.543; 95% CIs: 2.434 – 33.647 and p-value = 0.032; see Table S4 for full model estimates). With a yearly increase in storage time, plasma IGF-1 levels increased by 7.516 ng/ml, on average. Re-running our morphological and life-history analyses accounting for storage time effects on observed IGF-1 (by fitting storage time as a continuous covariate) revealed no qualitative differences in our main results.

We also investigated whether the time of day the blood samples were collected influenced plasma IGF-1 concentrations. In this dataset, all blood samples were collected sometime between 10am and 10pm with a mean sample collection time of 4pm (n=618 observations). We divided the distribution of sample collection times into tertiles namely, “Morning” for samples collected between 10am – 3pm; “Afternoon” for samples collected between 3pm – 6pm and “Evening” for samples collected between 6pm – 10pm. We then re-ran the IGF-1 model including sample collection time as a factor with three levels along with fixed effects of population density (continuous), sex (factor), litter size (factor) and maternal age at birth (linear and quadratic terms). All continuous predictors were z-standardized prior to model fitting. Like previous models, ELISA plate number, ELISA run date, maternal identity and birth year were fitted as random effects. Our analysis revealed no differences in plasma IGF-1 levels depending on the time of day the blood sample was collected (Figure S7; Table S5).

### Supplementary Figures

Figure S1: A schematic describing the relationships between IGF-1, proxies of resource availability, body mass and life-history traits. Proxies of resource availability can influence IGF-1 levels, body mass and life-history traits. IGF-1 can have a direct influence on morphological traits such as body mass. IGF-1 can also have a direct influence on life-history traits like survival and reproduction. In addition, IGF-1 could indirectly influence life-history traits through its effects on body mass.

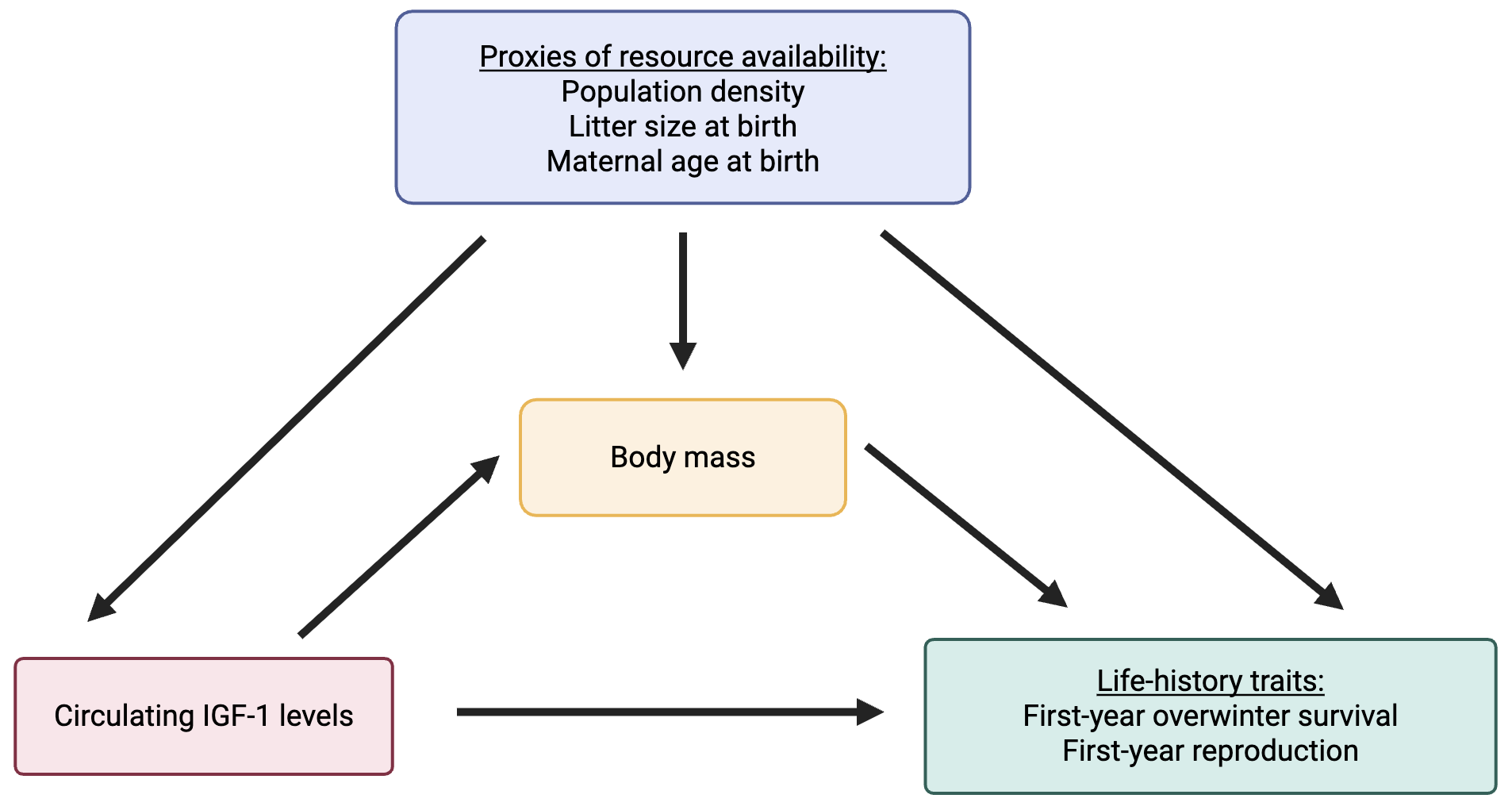

Figure S2: A) Histogram depicting the distribution of plasma IGF-1 levels in Soay lambs (n=669); B) Posterior distribution of the IGF-1 model (Table 1) with *y* representing observed values and *yrep* representing 200 draws from the posterior.

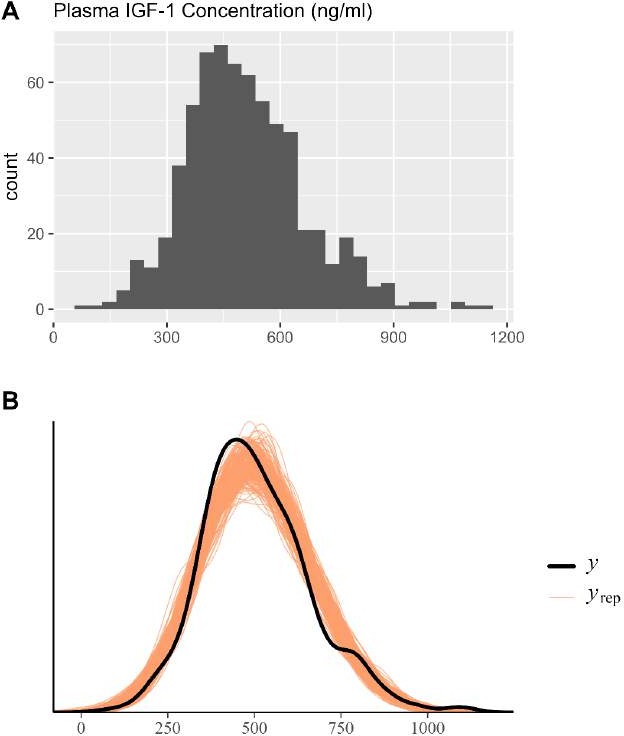

Figure S3: Posterior distributions of parameter estimates (medians and 95% credible intervals) of model investigating whether proxies of resource acquisition predict IGF-1 (Table 1; n=669 observations).

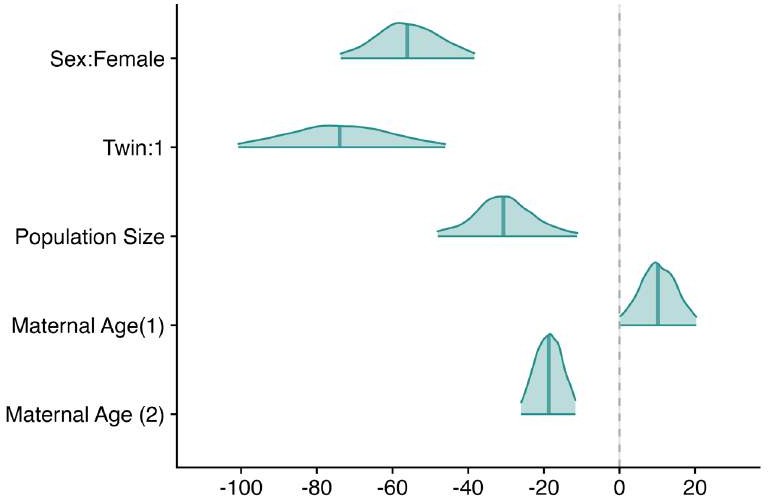

Figure S4: Posterior distributions of models investigating associations between August IGF-1 levels and A) August body mass; B) Post-natal growth; C) August foreleg length;

D) August horn length (in normal-horned individuals); E) First-year overwinter survival and F) First-year reproduction (Table S6, S7). Here, *y* represents observed values and *y_rep_* represents 200 draws from the posterior distribution.

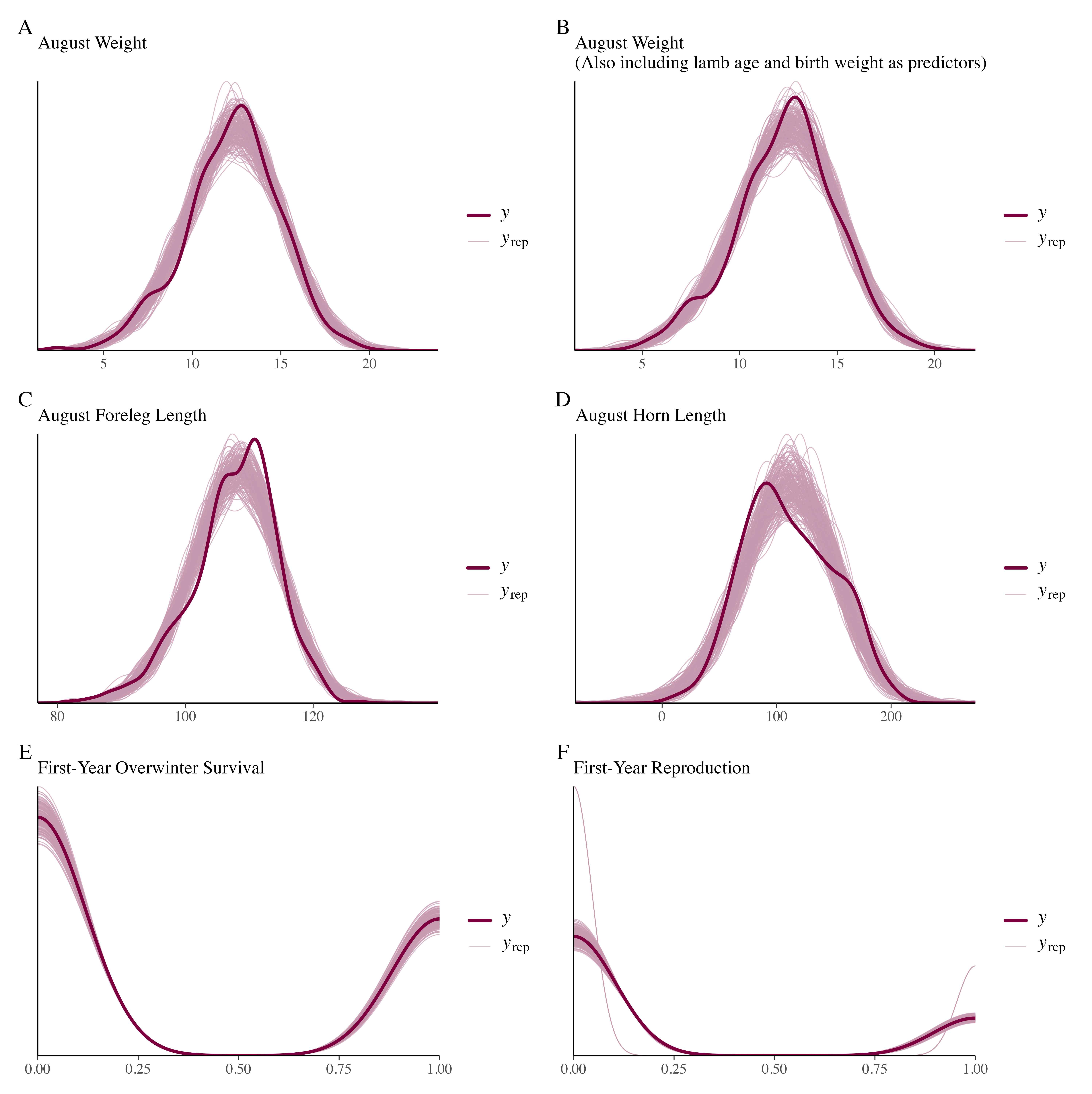

Figure S5: Association between plasma IGF-1 levels and different horn types (Model details in Table S3). Dots represent raw data and median and 95% CIs of posterior distribution displayed in black (n=669 observations).

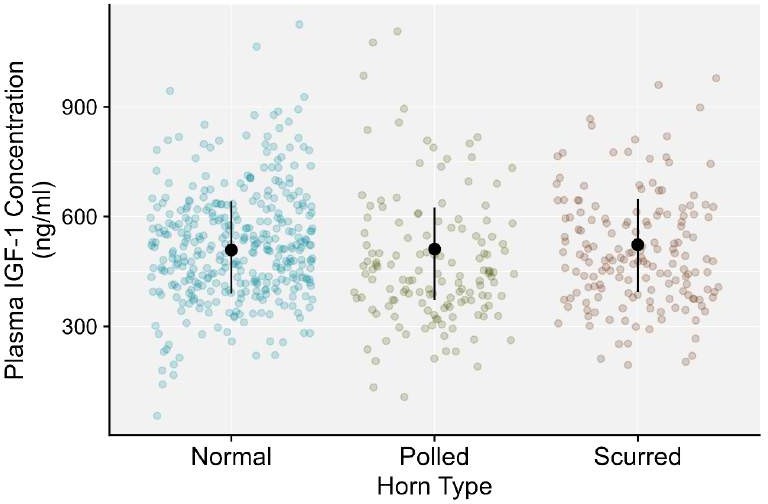

Figure S6: Association between plasma IGF-1 concentrations and sample storage time (in years). The dark line represents the median of the posterior distribution, and the lightly shaded lines are 500 draws from the posterior distribution. Points represent raw data (n=669 observations).

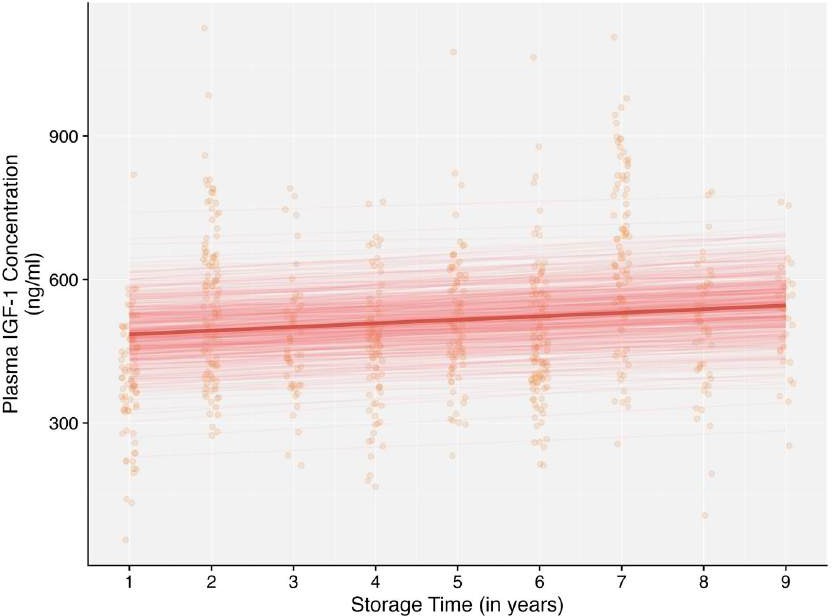

Figure S7: Association between plasma IGF-1 levels and the time-of-day blood samples were collected. Dots represent raw data with median and 95% CIs of posterior distribution displayed in black (n=669 observations).

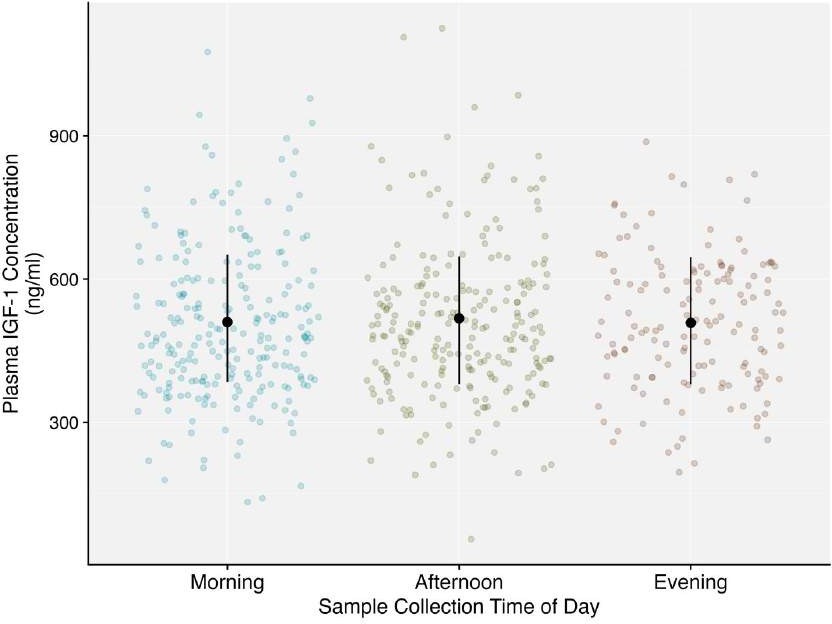

### Supplementary Tables

Table S1: Number of observations (or individuals) across different years, sexes, litter sizes, maternal ages and horn types.

| Year | Sex | Litter Size | Maternal Age | Horn Type |
| --- | --- | --- | --- | --- |
| 2014: 31 | Female: 337 | Singleton: 569 | Age 1: 7 | Normal: 362 |
| 2015: 46 | Male: 332 | Twin: 100 | Age 2: 47 | Polled: 141 |
| 2016: 89 |  |  | Age 3: 103 | Scurred: 164 |
| 2017: 110 |  |  | Age 4: 100 |  |
| 2018: 80 |  |  | Age 5: 81 |  |
| 2019: 83 |  |  | Age 6: 82 |  |
| 2020: 50 |  |  | Age 7: 67 |  |
| 2021: 101 |  |  | Age 8: 61 |  |
| 2022: 79 |  |  | Age 9: 46 |  |
|  |  |  | Age 10: 17 |  |
|  |  |  | Age 11: 8 |  |
|  |  |  | Age 12: 5 |  |
|  |  |  | Age 14: 1 |  |
|  |  | n=669 observations |  |  |

Table S2: Number of observations (or individuals) across different morphological and life-history traits.

| Trait | Number of observations |
| --- | --- |
| IGF-1 | 669 |
| August Body Mass | 666 |
| Post-Natal Growth | 545 |
| August Foreleg Length | 660 |
| August Horn Length | 361 |
| First-Year Reproduction | 656 |
| First-Year Overwinter Survival | 609 |

Table S3: Results from model investigating the association between plasma IGF-1 levels and sex, litter size, horn type, population density and maternal age (linear and quadratic terms) in Soay lambs (n=667; population size and maternal age at birth were standardized to mean=0 and standard deviation=1).

| Fixed Effect | Estimate [95% CI] | p-value |
| --- | --- | --- |
| Intercept | 568.320 | <0.001 |
|  | [441.000, 699.117] |  |
| **Sex: Female** | **-60.672** | **<0.001** |
|  | **[-84.342, -37.201]** |  |
| **Litter Size (Twin):1** | **-76.720** | **<0.001** |
|  | **[-103.416, -48.194]** |  |
| Horn Type: Polled | 2.777 | 0.861 |
|  | [-25.499, 33.107] |  |
| Horn Type: Scurred | 14.165 | 0.266 |
|  | [-10.432, 38.954] |  |
| **Population Size** | **-32.317** | **0.006** |
|  | **[-49.520, -13.085]** |  |
| **Maternal Age** | **10.335** | **0.036** |
|  | **[0.136, 20.007]** |  |
| **Maternal Age^2^** | **-19.071** | **<0.001** |
|  | **[-26.317, -11.799]** |  |
| Random Effect | Variance [95% CI] |  |
| Maternal Identity | 1934.665 |  |
|  | [828.303, 3366.747] |  |
| Birth Year | 173.439 |  |
|  | [0.000, 1477.282] |  |
| ELISA Run Date | 10181.120 |  |
|  | [159.580, 46898.660] |  |
| ELISA Plate Number | 3135.591 |  |
|  | [1139.377, 7126.428] |  |
| Residual | 11138.417 |  |
|  | [9718.963, 12734.603] |  |

Table S4: Results from model investigating the association between plasma IGF-1-1 levels and sex, litter size, storage time, population density and maternal age (linear and quadratic terms) in Soay lambs (n=669; storage time, population size and maternal age at birth were standardized to mean=0 and standard deviation=1).

| Fixed Effect | Estimate [95% CI] | p-value |
| --- | --- | --- |
| Intercept | 568.389 | <0.001 |
|  | [437.142, 686.518] |  |
| **Sex: Female** | **-56.180** | **<0.001** |
|  | **[-73.013, -39.769]** |  |
| **Litter Size (Twin):1** | **-71.833** | **<0.001** |
|  | **[-99.868, -45.661]** |  |
| **Storage Time** | **17.543** | **0.032** |
|  | **[2.434, 33.647]** |  |
| **Population Density** | **-22.134** | **0.012** |
|  | **[-37.630, -4.840]** |  |
| **Maternal Age** | **10.202** | **0.043** |
|  | **[0.096, 19.607]** |  |
| **Maternal Age^2^** | **-18.336** | **<0.001** |
|  | **[-25.889, -11.533]** |  |
| Random Effect | Variance [95% CI] |  |
| Maternal Identity | 2084.925 |  |
|  | [876.470, 3316.885] |  |
| Birth Year | 56.742 |  |
|  | [0.000, 720.399] |  |
| ELISA Run Date | 10056.520 |  |
|  | [250.901, 47218.330] |  |
| ELISA Plate Number | 2870.003 |  |
|  | [1008.537, 6470.085] |  |
| Residual | 11259.672 |  |
|  | [9844.370, 12906.369] |  |

Table S5: Results from model investigating the association between plasma IGF-1-1 levels and sex, litter size, sample collection time, population density and maternal age (linear and quadratic terms) in Soay lambs (n=618; population size and maternal age at birth were standardized to mean=0 and standard deviation=1).

| Fixed Effect | Estimate [95% CI] | p-value |
| --- | --- | --- |
| Intercept | 561.509 | <0.001 |
|  | [432.022, 689.184] |  |
| **Sex: Female** | **-55.468** | **<0.001** |
|  | **[-72.686, -37.784]** |  |
| **Litter Size (Twin):1** | **-78.592** | **<0.001** |
|  | **[-105.008, -50.607]** |  |
| Sample Collection Time: Afternoon | 8.308 | 0.464 |
|  | [-12.627, 30.023] |  |
| Sample Collection Time: Evening | -1.378 | 0.908 |
|  | [-25.316, 21.693] |  |
| **Population Size** | **-31.881** | **0.008** |
|  | **[-47.989, -14.617]** |  |
| **Maternal Age** | **13.414** | **0.011** |
|  | **[3.405, 23.229]** |  |
| **Maternal Age^2^** | **-18.030** | **<0.001** |
|  | **[-24.891, -10.361]** |  |
| Random Effect | Variance [95% CI] |  |
| Maternal Identity | 1400.632 |  |
|  | [102.637, 2682.257] |  |
| Birth Year | 138.812 |  |
|  | [0.000, 1302.499] |  |
| ELISA Run Date | 10093.750 |  |
|  | [1.004, 46714.190] |  |
| ELISA Plate Number | 3221.352 |  |
|  | [991.978, 7329.589] |  |
| Residual | 11008.332 |  |
|  | [9582.532, 12730.654] |  |

Table S6: Results from models investigating the association between plasma IGF-1 levels and

A) August body mass; B) Post-natal growth; C) August foreleg length; D) August horn length (in normal-horned individuals) in Soay lambs. Results from models of summer foreleg length and summer horn length controlling for proxies of resource availability (population density, litter size and maternal age), summer body mass, or both resource availability proxies and summer body mass, are also shown.

Median (95% CI)

| A) August Body Mass | 12.829 |  |
| --- | --- | --- |
| Intercept IGF-1  Sex (Female) | (12.171 – 13.481)  **1.057**  **(0.821 – 1.296)**  **-0.925**  **(-1.278 – -0.555)** |  |
| Litter Size (Twin) | - |  |
| Population Size | - |  |
| Maternal Age (1) | - |  |
| Maternal Age (2) | - |  |
| Variances |  |  |
| Maternal Identity Birth Year Residual | 1.493  (0.927 – 2.202)  0.572  (0.175 – 2.265)  4.109  (3.587 – 4.748) |  |
| B) Post-Natal Growth |  |  |
| Intercept IGF-1  Lamb Age (in days)  Birth Weight | 12.626  (12.001 – 13.282)  **1.173**  **(0.9779 – 1.362)**  **1.064**  **(0.923 – 1.211)**  **1.282**  **(1.133 – 1.428)** | 13.107  (12.55 – 13.636)  **1.028**  **(0.844 – 1.201)**  **0.96**  **(0.816 – 1.104)**  **1.074**  **(0.915 – 1.237)** |
| Sex (Female) | **-0.602**  **(-0.854 – -0.351)** | **-0.746**  **(-0.992 – -0.485)** |
| Litter Size (Twin) | - | **-1.114**  **(-1.563 – -0.659)** |
| Population Size | - | -0.241  (-0.789 – 0.325) |
| Maternal Age (1) | - | -0.041  (-0.193 – 0.106) |
| Maternal Age (2) | - | **-0.24**  **(-0.35 – -0.136)** |
| Variances |  |  |
| Maternal Identity | 0.434  (0.180 – 0.740) | 0.388  (0.140 – 0.662) |

|  | 0.615 | 0.402 |
| --- | --- | --- |
| Birth Year | (0.176 – 2.350) | (0.109 – 1.788) |
|  | 1.858 | 1.719 |
| Residual | (1.588 – 2.190) | (1.461 – 2.038) |

| C) August Foreleg Length |  | | | |
| --- | --- | --- | --- | --- |
| Intercept IGF-1  Sex (Female) Litter Size (Twin) Population Size Maternal Age (1)  Maternal Age (2) | 108.177  (106.882 – 109.416)  **3.294**  **(2.662 – 3.918)**  **-1.155**  **(-2.059 – -0.232)**  -  -  -  - | 107.146  (106.463 – 107.758)  **0.951**  **(0.535 – 1.355)**  **0.745**  **(0.162 – 1.294)** | 110.974  (109.338 – 112.595)  **2.128**  **(1.572 – 2.715)**  **-1.619**  **(-2.426 – -0.827)**  **-7.402**  **(-8.764 – -6.134)**  0.468  (-1.116 – 2.053)  **1.636**  **(1.180 – 2.095)**  **-1.544**  **(-1.887 – -1.2)** | 107.801  (107.076 – 108.494)  **0.891**  **(0.490 – 1.289)**  0.547  (-0.024 – 1.111)  **-1.922**  **(-2.890 – -0.963)**  0.215  (-0.308 – 0.725)  **0.671**  **(0.351 – 0.988)**  **-0.272**  **(-0.513 – -0.023)** |
| August Body Mass | - | **5.418**  **(5.072 – 5.739)** | - | **5.069**  **(4.704 – 5.437)** |
| Variances |  |  |  |  |
| Maternal Identity  Birth Year Residual | 5.963  (2.832 – 9.834)  1.893  (0.466 – 7.818)  29.921  (26.183 – 34.480) | 2.244  (0.984 – 3.865)  0.308  (0.003 – 1.784)  11.223  (9.728 – 12.964) | 7.332  (4.398 – 10.892)  3.531  (1.126 – 13.398)  21.245  (18.444 – 24.656) | 2.767  (1.419 – 4.393)  0.231  (0.002 – 1.596)  10.460  (9.094 – 12.192) |
| D) August Horn Length | 116.583 | 116.185 | 123.824 | 116.8 |
| Intercept | (109.012 – 123.389) | (111.169 – 121.300) | (115.637 – 132.128) | (111.107 – 122.456) |
| IGF-1  Sex (Female) Litter Size (Twin) Population Size Maternal Age (1)  Maternal Age (2)  August Body Mass | **11.854**  **(7.208 – 16.491)**  **-31.541**  **(-40.342 – -22.449)**  -  -  -  -  - | 1.432  (-3.240 – 6.462)  **-29.948**  **(-38.408 – -21.451)**  -  -  -  -  **19.320**  **(15.692 – 22.986)** | **8.786**  **(4.074 – 13.704)**  **-34.000**  **(-42.816 – -25.026)**  **-23.611**  **(-35.022 – -12.283)**  4.348  (-3.436 – 12.113)  **5.293**  **(1.486 – 9.318)**  **-3.558**  **(-6.33 – -0.479)**  - | 1.800  (-2.618 – 6.760)  **-30.364**  **(-38.406 – -22.639)**  -7.305  (-17.618 – 3.843)  3.051  (-1.589 – 8.159)  1.754  (-1.961 – 5.185)  0.575  (-2.128 – 3.524)  **18.615**  **(14.647 – 22.735)** |
| Variances |  |  |  |  |
| Maternal Identity  Birth Year Residual | 6.911  (0.015 – 175.377)  58.905  (2.502 – 253.988)  1165.198  (964.413 – 1366.982) | 14.063  (0.013 – 203.547)  16.695  (0.058 – 120.516)  883.338  (698.333 – 1045.488) | 8.084  (0.012 – 193.841)  60.343  (1.952 – 328.650)  1086.325  (885.634 – 1281.253) | 15.678  (0.029 – 194.525)  12.573  (0.047 – 124.041)  879.938  (698.281 – 1050.210) |

Table S7: Results from models investigating the association between plasma IGF-1 levels and

A) first-year reproduction and B) first-year overwinter survival in Soay lambs. Results from models controlling for summer body mass and both resource availability proxies and summer body mass are also shown.

Median (95% CI)

A) First-Year Reproduction

|  | -2.920 | -3.753 | -3.856 |
| --- | --- | --- | --- |
| Intercept | (-3.747 – -2.176) | (-4.566 – -3.022) | (-4.791 – -3.056) |
|  | **0.522** | 0.295 | 0.270 |
| IGF-1 | **(0.220 – 0.836)** | (-0.041 – 0.646) | (-0.110 – 0.630) |
|  | **2.579** | **3.518** | **3.556** |
| Sex (Female) | **(2.022 – 3.158)** | **(2.891 – 4.248)** | **(2.904 – 4.340)** |
|  |  |  | -0.157 |
| Litter Size (Twin) | - | - | (-1.074 – 0.721) |
|  |  |  | -0.361 |
| Population Size | - | - | (-0.847 – 0.106) |
|  |  |  | -0.087 |
| Maternal Age (1) | - | - | (-0.354 – 0.163) |
|  |  |  | 0.073 |
| Maternal Age (2) | - | - | (-0.161 – 0.296) |
|  |  | **1.189** | **1.242** |
| August Body Mass |  | **(0.881 – 1.540)** | **(0.868 – 1.641)** |
| Variances |  |  |  |
|  | 0.104 | 0.058 | 0.072 |
| Maternal Identity | (0.000 – 0.745) | (0.000 – 0.564) | (0.000 – 0.649) |
|  | 0.598 | 0.349 | 0.220 |
| Birth Year | (0.139 – 2.615) | (0.033 – 1.565) | (0.003 – 1.515) |
| B) First-Year Survival |  |  |  |
|  | -1.368 | -1.637 | -1.343 |
| Intercept | (-2.517 – -0.337) | (-2.856 – -0.462) | (-2.752 – -0.063) |
| IGF-1 | **0.410**  **(0.121 – 0.706)** | 0.119  (-0.210 – 0.440) | 0.107  (-0.251 – 0.448) |
|  | **0.941** | **1.274** | **1.247** |
| Sex (Female) | **(0.508 – 1.409)** | **(0.790 – 1.773)** | **(0.745 – 1.768)** |
|  |  |  | -0.425 |
| Litter Size (Twin) | - | - | (-1.225 – 0.378) |
|  |  |  | -0.706 |
| Population Size | - | - | (-2.057 – 0.505) |
|  |  |  | -0.131 |
| Maternal Age (1) | - | - | (-0.408 – 0.144) |
|  |  |  | **-0.291** |
| Maternal Age (2) | - | - | **(-0.565 – -0.042)** |
|  |  | **0.781** | **0.650** |
| August Body Mass | - | **(0.494 – 1.108)** | **(0.316 – 1.011)** |
| Variances |  |  |  |
|  | 0.852 | 1.000 | 1.170 |
| Maternal Identity | (0.187 – 1.982) | (0.194 – 2.353) | (0.297 – 2.692) |
| Birth Year | 2.014 | 2.286 | 2.286 |

(0.714 – 6.938)	(0.829 – 8.202)	(0.733 – 9.229)
